# Plasma FIB milling for the determination of structures *in situ*

**DOI:** 10.1101/2022.08.01.502333

**Authors:** Casper Berger, Maud Dumoux, Thomas Glen, Neville B.-y. Yee, John M. Mitchels, Zuzana Patáková, James H Naismith, Michael Grange

**Author notes:** These authors contributed equally (Casper Berger, Maud Dumoux, Thomas Glen).

## Abstract

Structural biology inside cells and tissues requires methods able to thin vitrified specimens to electron transparent thicknesses. Until now, focused ions beams based on gallium have been used. However, ion implantation, changes to surface chemistry and an inability to access high currents limit Gallium as an ion beam source. Here, we show that plasma-coupled ion sources can produce cryogenic lamella of vitrified human cells in a robust and automated manner, with quality sufficient for pseudo-atomic structure determination. In addition, these lamellae were produced in a prototype microscope equipped for long cryogenic run times (>1 week) and with multi-specimen support fully compatible with modern-day transmission electron microscopes. We demonstrate for the first time that plasma ion sources can be used for structural biology within cells, determining a structure in-situ to 4.9 Å and describing a workflow upon which different plasmas can be examined to streamline lamella fabrication further.

## Introduction

Cryo-electron tomography (cryoET) enables the structural study of macromolecular complexes within their native cellular environment. Understanding the structural landscape of macromolecules in relation to their subcellular environment is key in linking structure to function. Since most biological samples are too thick to directly image with a transmission electron microscope (TEM), methods to thin down biological material to the < 300 nm necessary for electron transparency have been instrumental for enabling cryoET of the internal regions of cells. Focused ion beam (FIB) milling was originally developed for material science applications, such as microchip manufacture, at room temperature, where its ability to precisely shape objects on the nanoscale revolutionised electronics^1,2^. More recently, the approach has been adapted for use with biological samples through the adaptation of existing dual beam FIB/scanning electron microscopes (FIB/SEM) with cryogenic stages enabling frozen-hydrated samples to be shaped and thinned to thicknesses enabling subsequent cryoET analyses^3,4^. This capability has enabled intracellular structural studies on a wide variety of biological samples and to pseudo-atomic resolution^5–9^.

Current FIB/SEM instruments used to prepare FIB lamella are not optimised for cryogenic applications^10–12^, reducing their throughput. The large sample chamber (~40 L), originally designed for 8” Si wafers at room temperature, has a vacuum typically of the order 10^−6^ mbar, dropping by a ~10-fold when cold. This results in ice (re)deposition, with rates of up to 85 nm/h reported^11,13^. This limits number of lamellae that can be prepared on one grid in an experiment and impacts the proportion of successful lamella^11^. Studies have recently demonstrated success with automated FIB-lamella fabrication^11,12,14,15^, but use of gas cooled systems limits length of overnight runs and ice growth effectively limits the number of lamella that can be prepared in one automated session.

Lamella fabrication to-date for structural biology has typically been performed with a gallium FIB. Gallium works well as a metal ion source due its low melting point, low volatility, low vapour pressure, good emission characteristics and good vacuum properties^16^. Gallium also sputters a range of different materials efficiently. The point source nature of a gallium ion source enables the beam to be tightly focussed^17^ enabling control at the tens to hundreds of nanometer scale. At higher beam currents the beam becomes increasingly divergent, limiting the bulk milling these beams are capable of. Implantation is likely from any focussed ion beam^18^, but gallium has been shown to react with various samples, migrate and significantly alter the original structure^18–20^. Alternative approaches are therefore of interest to reduce these effects.

An alternative is inductively coupled plasmas generated from a gas, e.g., nitrogen, oxygen, xenon, or argon. These plasma sources are comparable in size to gallium at low current but have smaller probe sizes than gallium at larger beam currents^21^. This means that in principle plasma beams are highly suited to milling large volumes but remain capable of the fine milling required for thinner, more fragile, lamella used in life sciences^22^.

Here, we describe a protocol for automated plasma FIB milling lamella of cryogenic cellular samples. We were able to hold multiple specimens at cryogenic temperatures at low contamination rates (<2 nm / hr) for weeks. We characterised the milling rates of plasma ion sources (O, N, Ar, Xe) at a range of currents. We demonstrate that an argon plasma source can produce lamellae with a high success rate (~80%). The lamellae were used to generate a cryoET dataset of hundreds of tomograms, which show thicknesses, features, and resolution on a sub-nanometre scale, enabling sub volume averaging of the human 80S ribosome to a resolution of ~4.9 Å, with the well-ordered regions of the structure at resolutions close to the Nyquist limit of 3.8 Å. Our study demonstrates for the first time the use of a plasma ion source to enable structural biology *in situ* and paves a way for higher throughput.

## Results

### A plasma FIB/SEM for screening and imaging multiple samples with minimal contamination over long time scales

We used a custom designed plasma FIB/SEM microscope (Supplementary Fig. 1) equipped with a coincident FIB and SEM inside a chamber with redeposition rates < 2 nm/hr (Supplementary Fig. 2), and a stage with rotational freedom of +14° to −190°. The sample chamber vacuum is maintained at a pressure of ~1 × 10^−7^ mbar. The redesigned sample chamber is approximately 6 litres in volume. Up to 12 clipped grids can be mounted into a multi-specimen cassette which can be robotically loaded into the chamber, one at a time (Supplementary Fig. 1). The PFIB can be configured to be used with different ion sources, including xenon, oxygen, argon. The cryo box (anti-contaminator) and stage are braid cooled from liquid nitrogen dewars that are filled automatically when needed, allowing continuous cold runtimes of up to a week or more. The characteristics of this microscope, including minimal ice growth (Supplementary Fig. 2), robotic sample transfers and long cryo runtimes enabled preparation of FIB-lamellae on biological specimens for up to one week. In effect the number of lamellae that can be produced is sample dependent. The high-vacuum exchange of samples and robotic sample entry meant that samples could be handled with minimal damage or disruption.

### Milling rates of plasma ions on vitreous cellular samples

Determining the sputter rate of each plasma beam can inform the design of milling protocols. Others have already characterised the sputter rate for different materials^23^, but not on vitrified biological samples. We used a second microscope (Thermo Scientific Helios Hydra with gas-cooled cryogenic stage) with the same ion column as the first to determine sputter rates more accurately for xenon, nitrogen, oxygen, and argon, at normal incident angles. Based on 3 experimental repeats on different regions of the same plunge frozen yeast sample, our measurements show that for vitrified biological samples xenon has the highest sputter rate of 16.8 ± 0.2 µm^3^/nC. Nitrogen and oxygen have values of 10.6 ± 0.2 and 10.0 ± 0.4 µm^3^/nC respectively, while argon is 4.3 ± 0.21 µm^3^/nC (Supplementary Fig. 3; Table 1). This compares with a sputter rate of 7.7 µm^3^/nC for gallium in ice^24^. These sputter rates are ~20-100 times higher than those reported at 30kV in Silicon for these plasma beams^23^. This is of the same order as the factor of ten that is suggested within the community as a rule of thumb for the difference between milling in vitrified ice and silicon^25^. During lamella fabrication for cryoET, typical FIB angles for milling can range between 8 – 13°, with angles up to ~25° being used for some applications. We measured the sputter rate of vitrified samples using xenon and argon at currents between 20 pA up to 2 nA (currents typically used for lamella fabrication^13,26^ at angles from 10° to 40°, and calculated sputter rates. (Supplementary Fig. 3; Table 2)). Our results show that at high angles, there was a greater difference in sputter rates between the two gases (at 90° xenon is 3.7x greater) compared to shallow angles (10° xenon is 2.7x greater), suggesting that the incidence angle influences sputter efficiency in vitrified water, and that this dependence varies with milling gas.

**Table 1.** Measured sputter rates taken from triplicate measurements on plunge-frozen yeast samples. Sputter rates for Xenon, Nitrogen, Oxygen and Argon are shown. Sputter rates were measures as described in Supplementary Fig. 3. Errors shown are the standard error derived using the least squares method from a linear line of best fit that passes through the origin.

| Plasma | Xenon<br>(MW 131.29) | Nitrogen (MW<br>14) | Oxygen (MW<br>15.99) | Argon<br>(MW 39.95) |
| --- | --- | --- | --- | --- |
| Sputter rate<br>($\mu\text{m}^3/\text{nC}$) | $16.7 \pm 0.2$ | $10.6 \pm 0.2$ | $10.0 \pm 0.4$ | $4.3 \pm 0.1$ |

**Table 2.**
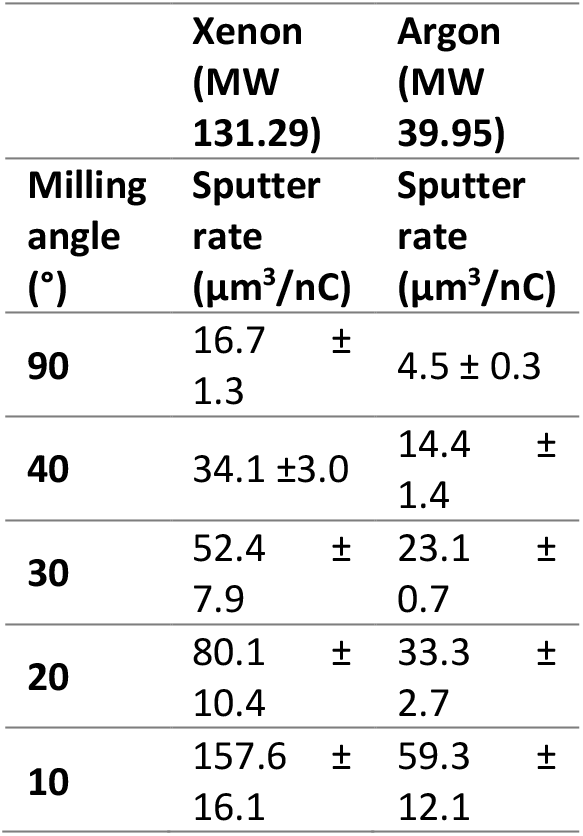
Measured sputter rates taken from triplicate measurements on plunge-frozen HeLa samples with varying milling angles. Sputter rates for Xenon, and Argon are shown. Sputter rates were measures as described in Supplementary Fig. 3. Errors shown are the standard error derived using the least squares method from a linear line of best fit that passes through the origin.

|  | Xenon<br>(MW<br>131.29) | Argon<br>(MW<br>39.95) |
| --- | --- | --- |
| Milling<br>angle<br>(°) | Sputter<br>rate<br>($\mu\text{m}^3/\text{nC}$) | Sputter<br>rate<br>($\mu\text{m}^3/\text{nC}$) |
| 90 | 16.7 ± 1.3 | 4.5 ± 0.3 |
| 40 | 34.1 ± 3.0 | 14.4 ± 1.4 |
| 30 | 52.4 ± 7.9 | 23.1 ± 0.7 |
| 20 | 80.1 ± 10.4 | 33.3 ± 2.7 |
| 10 | 157.6 ± 16.1 | 59.3 ± 12.1 |

### Plasma is a suitable ion source for automated, high throughput lamella fabrication

While xenon has the greater sputter rate, we opted to use argon for our first experiment of fully automated lamella fabrication. This is because we had accumulated more experience in the control of argon plasma to produce smooth flat surfaces. We reasoned that a smoother surface would correlate with lower damage and therefore a better initial test of the approach. Since the instrument is not limited by ice redeposition in the chamber for these initial experiments the additional time taken by argon was not critical. Over two days 34 lamella were milled automatically from two grids (Fig. 1). The average time taken for milling was approximately 45 mins per lamella; 32 minutes of coarse milling and 13 minutes of fine milling (details in Table 3).

**Table 3.** Milling parameters used for lamella preparation.

| Step | Milling<br>current<br>(pA) | Width<br>($\mu\text{m}$) | Height<br>($\mu\text{m}$) /<br>Overlap<br>(%) | Offset<br>from<br>lamella<br>(nm) | Milling<br>depth<br>($\mu\text{m}$) | Drift<br>correction<br>interval<br>(s) <sup>642</sup> |
| --- | --- | --- | --- | --- | --- | --- |
| Rough<br>milling | 2000 | 15 (top)<br>14<br>(bottom) | 7 (top)<br>7<br>(bottom) | 1500 | 1.875 | 500 |
| Medium<br>milling | 200 | 13.3<br>(top)<br>13<br>(bottom) | 200% | 600 | 1.41 | 500 |
| Fine<br>milling | 60 | 12.7<br>(top)<br>12.2<br>(bottom) | 200% | 300 | 1.125 | 60 |
| Polish 1 | 60 | 12 | 200% | 150 | 0.5625 | 30 |
| Polish 2 | 20 | 12 | 200% | 0 | 0.5625 | 10 |

**Figure 1.**
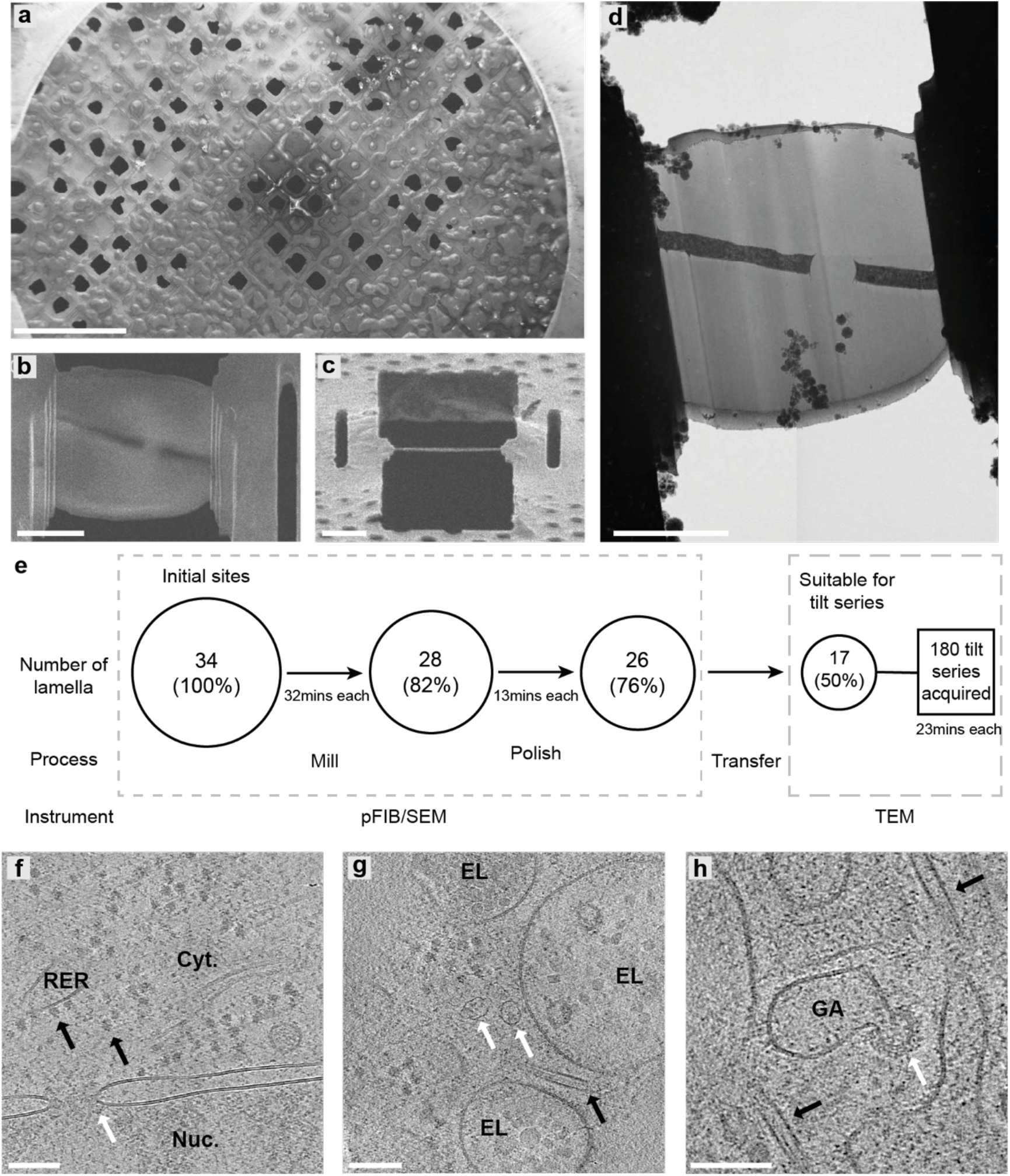
Overview of lamella production using plasma FIB milling. **(a)** SEM overview of the uncoated grid. **(b)** SEM image of a finished lamella and **(c)** its respective FIB image. **(d)** Low magnification TEM image of the same lamella. Scale bar are 500 µm in **(a)** and 5 µm in **(b), (c)** and **(d)**. The chart in **(e**) shows the number of successful lamella sites at each stage with the time taken between coarse and finer milling steps, starting with 34 initial sites down to 17 on which tilt series were acquired. **(f-h)** Representative tomographic slices containing several sub-cellular features, including **(f)** nucleus (Nuc.) and cytosol (Cyt.) with a nuclear pore (white arrow) in the nuclear envelope. Ribosomes (black arrows) are visible both free within the cytosol and tethered to the rough-endoplasmic reticulum (RER). scalebar: 100 nm; **(g)** endo-lysosomal vesicles (EL), a microtubule (black arrow) and vault complexes (white arrows). Scale bar: 100 nm; **(h)** microtubules (black arrows) and vesicle budding from the Golgi (GA) via a retromer coat (white arrow). The full tomographic volume is shown in Supplementary Video 1. Scale bar: 100 nm.

Once completed the grids were transferred to a compatible TEM in the same cassette, preserving the milling angle close to perpendicular to the tilt axis (robotic automation still leads to some slight rotational variability upon loading), eliminating need for manual handling. No lamellae were lost during sample transfer between PFIB and TEM.

From the 34 positions initially prepared, 26 lamellae were imaged in the TEM (i.e., were thin enough to enable transmission electron imaging), giving a successful fabrication rate of 76% (Fig. 1e). Of those, 17 were deemed suitable for tilt series acquisition (excluding lamella with incomplete vitrification, cracks, or unsuitable acquisition areas (e.g nucleus)). Therefore, of the lamella produced 65% (50% of initial set-up positions) were able to produce data from electron cryo-tomography; a total of 180 tilt-series.

### Electron cryo-tomography of plasma FIB milled lamella shows vitreous ice and allows an assessment of cellular morphology and ultrastructure

Tomograms acquired from the lamellae generally exhibited characteristics associated with vitrified cellular tomograms (no Bragg reflections within images, no aggregation-segregation induced contrast, uniform rounded membranes). We could observe a wide variety of macromolecular complexes (Fig. 1f-h, Supplementary Video 1) in the reconstructed tomograms including nuclear pore complexes^27,28^, microtubules^29^, vault complexes^30^, and vesicle budding via retromer coat proteins^31^. The reconstructed tomograms had an overall mean thickness of 250 nm ± 70 nm for the recorded tilt-series (Fig. 2). Although manually prepared FIB-lamellae are generally thinner^32^, the lamellae generated here were suitable for successful TEM tilt-series acquisition, observing cellular features in tomograms > 350 nm thick (Fig. 2).

**Figure 2.**
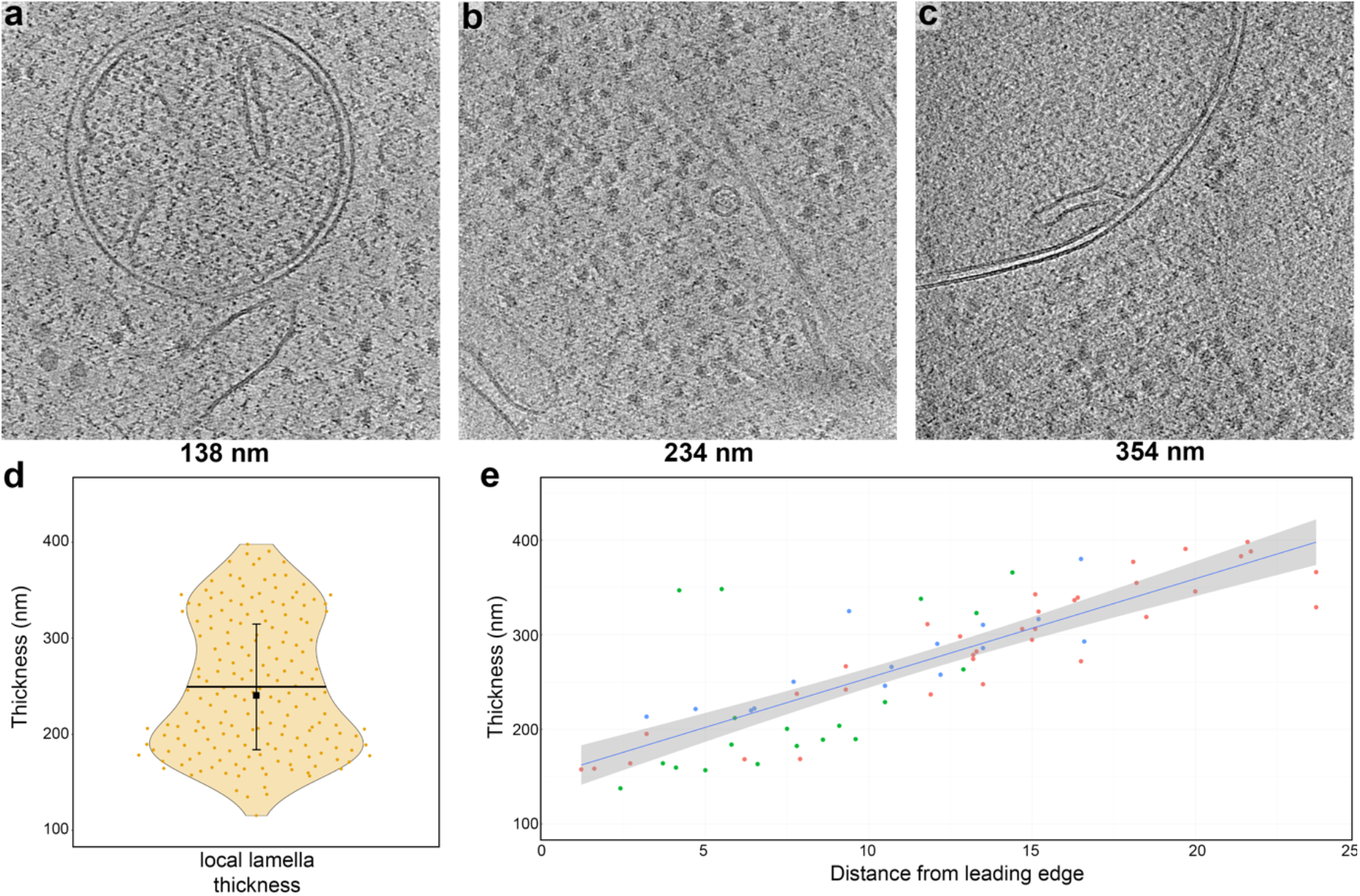
Plasma-milling produces lamellae with a range of thicknesses that exhibit usable sub-cellular tomographic data. (a-c) representative tomographic slices of tomograms recorded on lamellae with different thicknesses (indicated below each panel). Scale bars: 100 nm. (d) Violin plot showing the distribution of tomogram thickness as determined in the reconstructed tomograms in all 180 tilt-series (orange, mean SD; 250 nm ± 70). Horizontal black bars indicate the mean thickness, error bars the standard deviation and the black square the median (241 nm). (e) Scatter plot for tomogram thickness and the distance to the front of the lamellae for all tomograms recorded on three different lamellae (datapoints coloured in green, red and blue for each lamella). A linear trend line (blue) is shown, with the 0.95 confidence interval shown in grey. Plots for 8 individual tomograms are also shown in Fig. S5.

We measured the thickness of all lamellae and the distance from the leading edge at which the tomogram was acquired (Fig. 2; Supplementary Fig. 4). Interestingly, a number of lamella showed a gradient of thickness^13^, with some lamella being twice as thick at the back of the lamella as toward the front (<200 nm vs >350 nm), while others were relatively flat. The lamella with the greatest thickness gradient were the longest (Supplementary Fig. 4). There is a need to optimise the protocol to best reflect the cellular size/morphology to achieve flat lamella. In the current test case, no over tilt was used in coarse or fine milling steps to attempt to optimise flatness^13^.

### Sub-volume averaging at sub-nm and the effect of particles at the lamella edge

We determined structures of the human ribosome within the HeLa cells. We could obtain a structure of the human 80S ribosome using 15628 particles from 17 lamellae with a global resolution of 4.9 Å, and local resolution of 3.8 Å for large areas of the 60S subunit (Fig. 3, Supplementary Fig. 5, Supplementary Movie 2 and 3). During refinement of the structure to sub-nm resolutions, the 60S and 40S diverged in resolution, with the 60S approaching a resolution close to the Nyquist information limit (3.8 Å), while the 40S was a resolution above 8 Å. As the particle population is taken from non-arrested HeLa cells, our data captures the conformational states of the ribosome during translation. These data are available on EMPIAR (EMPIAR-XXXXX) should a study wish to probe human ribosome dynamics further.

**Figure 3.**
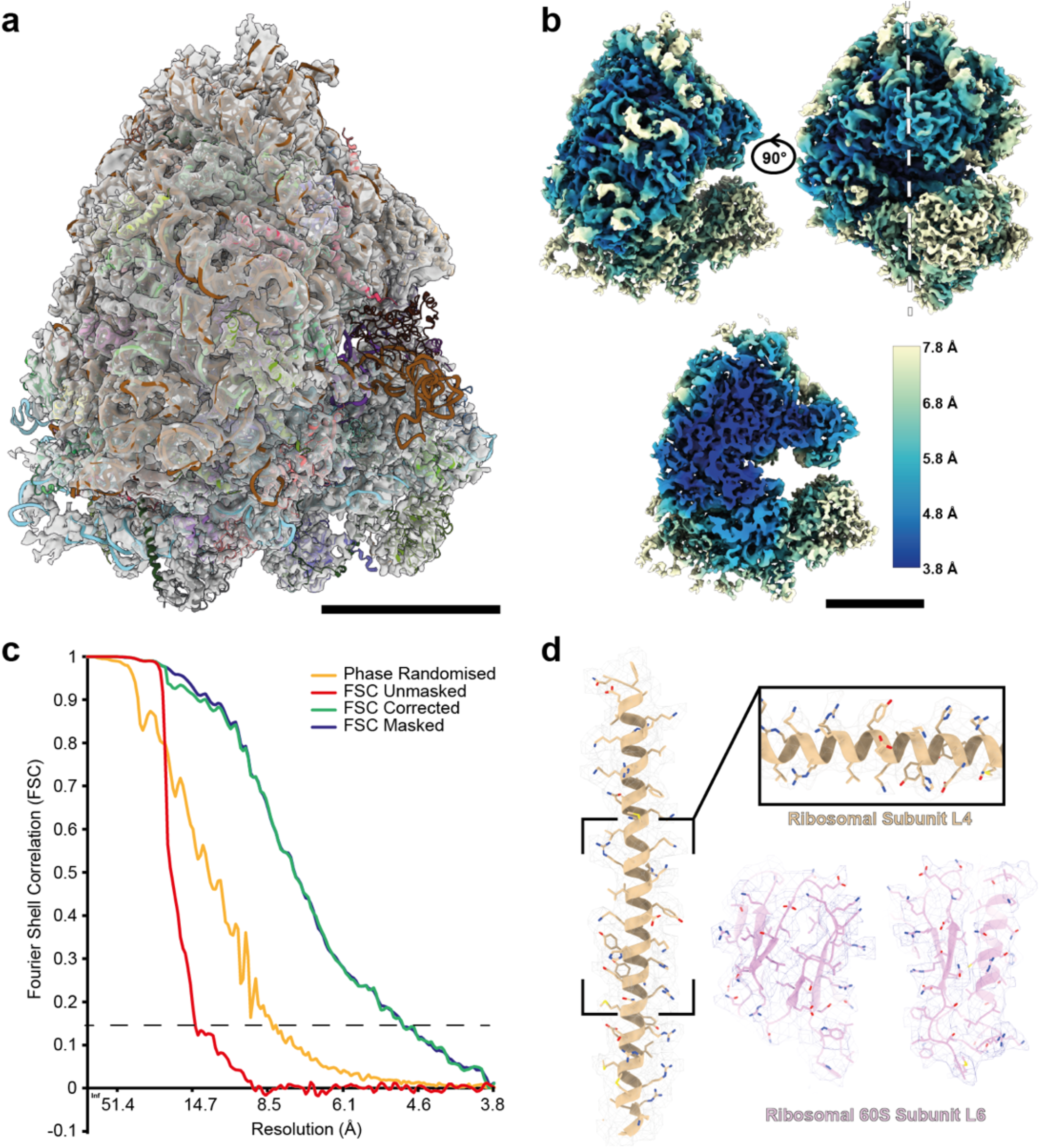
Structure of the human 80S ribosome obtained from cellular tomograms. **(a)** Density map of the 80S ribosome with fitted single-particle determined model of the human ribosome obtained from isolated ribosomes (PDB: 4UG0^46^) scale bar: 10 nm. **(b)** The same density map colour-coded for the local resolution from the outside (top) and a central slice (bottom, white line) (scale bar: 10 nm). **(c)** Fourier shell correlation for the 80S ribosome gives a masked global resolution of 4.9 Å (dotted line, FSC 0.143). **(d)** Representative fits into the EM density for different ribosomal regions (60S subunits L4 (gold) and L6 (magenta), suggesting quality of the map.

We interrogated the effect of plasma on damage propensity close to the milling boundary of the lamella. Currently it is difficult to ascertain the damage induced by an ion beam during lamella fabrication for life science samples; diffraction methods are not possible on cellular samples and sub-volume averaging has, until recently^8,33^, been limited to low resolution. The ability to produce structures at sub-nm resolutions offers the opportunity to determine damage due to ion beam milling (based on ability to align particles to a given resolution). Boundary models for all 180 tomograms were produced and then interpolated to form a geometric model of the tomograms. This enabled robust modelling of the lamella geometry local to each tomogram (Supplementary Fig. 6). When we compared the number of particles 3D classification with the number after picking, there was an appreciable loss of particles within 20 nm from the lamella edge (Supplementary Fig. 7). The particle positions taken from sub-volume averaging of 3D classified 18,119 particles were then used to calculate the distance at which each ribosome was located relative to the boundary of the lamella/tomogram. Across the 180 tomograms, 2099 ribosomes were calculated to be within 30 nm of the lamella edge (mean tomogram thickness 190 ± 39 nm). 2099 particles outside of this 30 nm layer were then randomly taken as a comparison (mean thickness 216 ± 49 nm). Each set of particles were then independently subjected to sub-volume averaging using a new global 3D refinement. Interestingly, particles taken from the 30 nm layer refined to a resolution of 9.2 Å compared to a resolution of 8.0 Å for the ribosomes outside of that distance (Fig. 4; Supplementary Fig. 8). The B-factors^34^ for the two particle populations also suggest particles within the 30 nm are of lower reconstruction quality, with a 24 % difference between the two particle populations (628 Å^2^ vs 480 Å^2^) (Fig.4).

**Figure 4.**
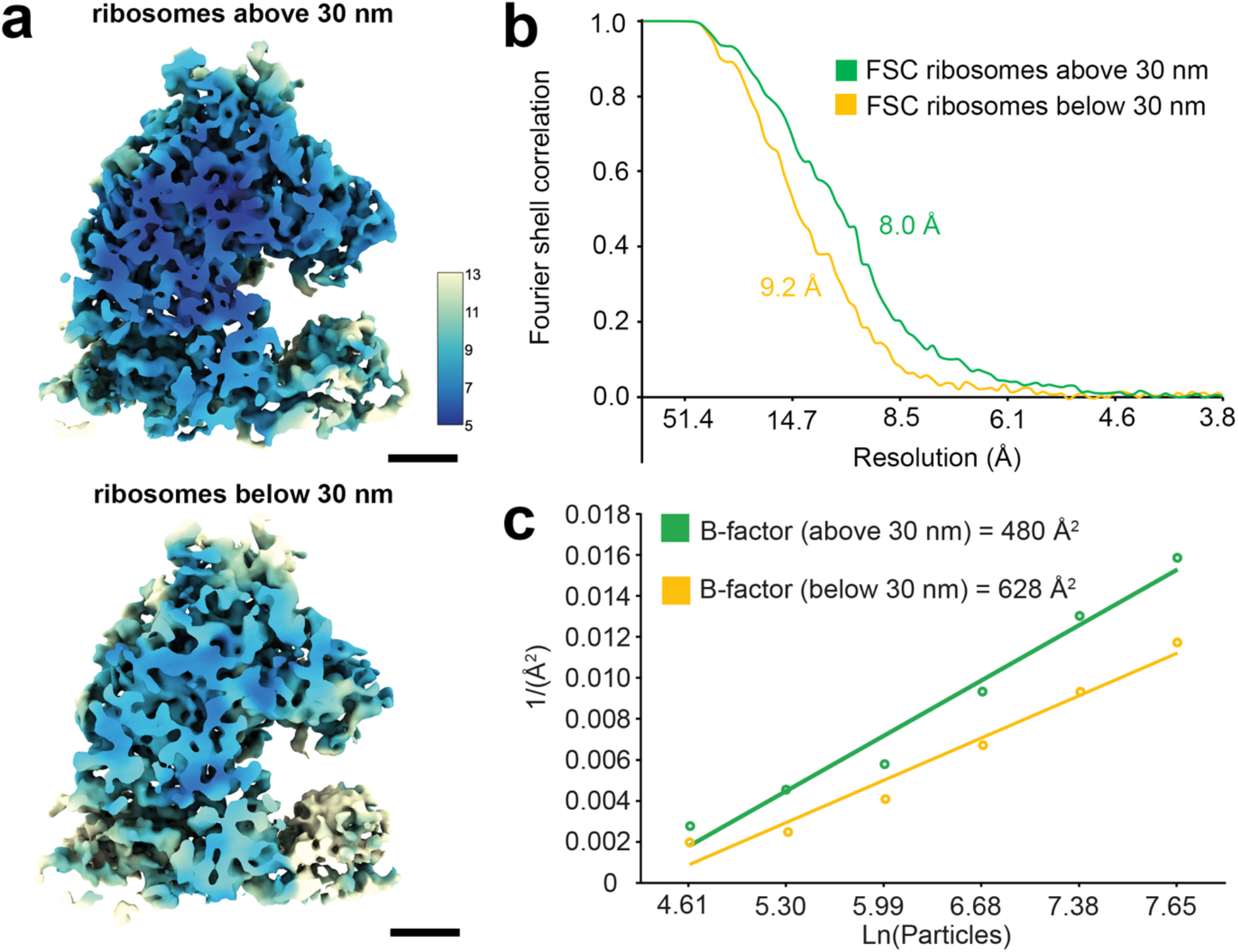
Effect of distance from the lamella edge on the ability to determine ribosome structures. **(a)** Ribosome structures were determined from acquired tomograms, where 2099 particles taken from outside **(top)** and within **(bottom)** the 30 nm of the edge of the lamella. Ribosomes are shown colour by the local resolution of the maps (key in Å). Scale bars are 5 nm and 1 nm (inset). **(b)** FSC curves are shown for the two maps, with a global resolution of 9.2 Å for the ribosomes within 30 nm of the edge (yellow) and 8 Å for ribosomes outside of this distance (green). **(c)** B-factors determined for particles above (green) and below (yellow) the 30 nm distance from the lamella edge after sub-volume averaging. These are calculated as 2 over the slope (Å^2^).

## Discussion

The determined sputter rates for each plasma (Table 2; Supplementary Fig. 3) are non-linear with molecular weight (MW) of the used gas. While xenon (MW 131.29) has the greatest sputter rate and largest MW, argon (MW 39.95) has the lowest sputter rate while weighing more than nitrogen (MW 14) and oxygen (MW 15.99). The sputter rates for oxygen and nitrogen are the same within error. While there is an expectation that a greater MW should correlate with sputter rate from first logic, the sputter rate for different plasmas has been demonstrated previously to be material dependent^23^. In our case, the values are calculated for (vitreous) biological samples. Interestingly, the angle dependence we were able to calculate for argon and xenon shows that closer to the glancing angle (smaller milling angles) the sputter rates increase greatly. However, more surprisingly the sputter yield for argon becomes a much greater relative yield with respect to xenon, with the relative sputter rates changing from 4.5 µm^3^/nC to 59.3 µm^3^/nC from 90° incidence to 10° incidence, suggesting a high angular dependence on sputter rates. Interestingly, this effect is greater for argon than for xenon (13-fold increase for argon vs 9.4-fold increase for xenon). These observations are consistent with published simulations^35^ but mean that at very shallow angles the choice of gas becomes less critical (between Xe and Ar). Other contributory factors, such as curtaining propensity (either based on probe characteristic or defocus variation) should therefore be considered to a greater extent at these angles.

The mean free path of an electron at 300 kV in water is approximately 300 nm^36,37^. This limits the thickness of samples that can be imaged by electrons. Traditionally lamella below 300 nm are prepared to achieve sufficient contrast for cryoET; a simple rule of thumb is that the thinner the lamella the better the contrast. Although higher in contrast, such thin lamellae (< 150 nm) suffer from having a greater proportion of the volume subject to milling induced effects and images a smaller volume of biological space reducing contextual information. An open question is whether plasma FIB can be used autonomously to fabricate lamella suitable for *in situ* structural biology. We report a protocol that produces lamella with a mean thickness of 250 ± 70 nm that runs unsupervised. We show that the lamella produced in this way yield information rich tomograms. From these tomograms it was possible to use sub tomographic averaging to obtain a structure of the human ribosome to 4.9 Å within lamellae. This compares extremely well with the current contemporary studies in cryo-tomography; ribosomes in isolated bacteria to 3.5 Å^33^, ribosomes from lamella of *S. Cerevisiae* (Yeast) to a resolution of 5.1 Å^38^ and recently lamella of isolated vertebrate myofibrils where the thin filament was determined to 4.5 Å resolution^8^. The milling process was complete in 2 days and the TEM data acquisition in 4 days. Thus, at a practical level, the protocol opens a route to rapid *in situ* structural biology with resolutions at or exceeding the highest currently available.

We can only speculate why such high-quality data were obtained from lamella that would be regarded as “thick” (i.e. in comparison to a gallium milled example of ~185 nm^32^). Any implantation of the milling ion into lamellae during fabrication will increase the proportion of inelastic vs elastic scattering reducing image contrast. Argon has over 1/3 less electrons than gallium thus for an equivalent level of implantation argon would be less distorting due to lower local change in potential derived from the implanted ion species. Accurate measurement of gallium and argon implantion upon vitrified samples requires scanning electron methods and/or electron energy loss spectrometry that are beyond the scope of this work. Notably both these approaches require very large electron fluences (2000-3000 e^−^/Å^2^) which would severely damage the sample. Secondly, the very high vacuum in the both the FIB/SEM and the TEM instrument used here have practically eliminated redeposited ice contamination in the chamber. Thus, the milled lamella thickness that we fabricate is virtually the same thickness of the lamella that is measured during the TEM experiment. This is in contrast to other approaches where the lamella that is measured by TEM is often the milled thickness plus a layer of (contrast reducing) redeposited material^11^. The transfer between the autoloading devices on the machines employs a portable cassette filled with liquid nitrogen at atmospheric pressure, some ice contamination was seen on some of our early lamella (Fig. 1). Subsequently using careful precautions, (low humidity, heating and drying) of all equipment and freshly (< 2 hrs) dispensed liquid nitrogen we have been able greatly reduce ice contamination. Vacuum transfer between microscopes would significantly simplify the process, eliminate ice contamination, and allow a more flexible workflow.

A robust determination on the effect of FIB milling on the damage caused to the surface of the lamella (structural changes (amorphisation if lattice) rather than vitrification) is yet to be reported. Ribosomes populate much of the cell making them an attractive model to assess the effect of milling. We were able to show via sub-volume averaging, that particles near the lamella edge contain less high-spatial frequency information compared to particles further away from the edge. In the 30 nm closest to the top or bottom surface of the lamella, averaging the ribosomes yields the resolution of 9.2 Å vs 8.0 Å for those particles found in the middle of the lamella. In addition, the particles producing the higher resolution structure were taken from tomograms with a greater mean thickness (190 nm vs 216 nm); this suggest particles in the central region of lamellae are more useful for sub-volume averaging despite the reduction in signal-to-noise due to this greater thickness. This was also evident from the calculated B-factors. Taken together, these data suggest that argon plasma, even at low currents, leads to some structural change within the first 30 nm of the lamella edge. Since the diameter of the ribosome is 25 Å, a portion of the damage could result from partial ablation of the ribosome or ribosomes immediately at the surface. Nevertheless, this is the first empirical measure of (plasma) beam damage for biological samples, and this analysis establishes a standard that can be used by us and others to robustly evaluate other milling protocols and plasma chemistries.

## Conclusions

Electron cryo-tomography is the frontier of structural biology but there is a pressing need to streamline workflows to enable its widespread adoption. Here we present a robust method for lamella fabrication using plasma FIB. This is the first demonstration of plasma for usage in lamella fabrication for cryoET. Our results show that a plasma cryo FIB/SEM with low contamination rates and with a robotic multi-specimen loading device can streamline the fabrication of lamella for pseudo-atomic structure determination. In our first attempt using argon plasma, we produced around 20 high quality lamella per day. We have accurately quantified the depth of damage from plasma milling with argon but show this does not prevent pseudo atomic resolution structure determination. We conclude that plasma FIB lamella fabrication is a suitable route for high throughput *in situ* structural biology.

## Materials and Methods

### Dual Beam FIB/SEM Descriptions

Milled samples and plasma characteristics data were acquired on either i.) a dual-beam focused ion beam scanning electron (FIB/SEM) “Helios Hydra” microscope (ThermoFisher Scientific, Oregon, USA) equipped with a cryogenic stage and plasma multi-ion source (argon, nitrogen, xenon, and oxygen) or ii.) a dual-beam focused ion beam scanning electron microscope with redesigned sample chamber and loading mechanism (ThermoFisher Scientific, Oregon, USA). Briefly, this microscope was equipped with several modifications to enable plasma FIB milling within a small, enclosed, (ultra) high vacuum chamber.

#### Chamber/Autoloader/Stage

The chamber volume of the system is much smaller than the Helios Hydra, with a reduction in volume from approximately ~0.04 m^3^ to < 0.01 m^3^ (less than a 5th of the volume of a conventional Helios-type chamber). This enables a vacuum on the order of 1 × 10^−7^ mbar to be achieved. It includes ports for the plasma FIB and SEM columns, cryogenic stage, robotic multi-specimen entry/exit (Autoloader), charge dissipation via a platinum metal sputter target and organometallic platinum protective coating chemistry via a gas injection system (GIS). Its volume reduces the contamination rates of a conventional chamber to < 2 nm/hr. The chamber also comprises of a stage which allows loading of grids assembled into compatible mounts (autogrids) via the Autoloader. The autogrids are also compatible with the Titan Krios for imaging via TEM and can be shuttled between the instruments via a cassette without need for manual handling. Up to 12 samples may be stored and recalled from the autoloader to the main stage and then subsequently transferred to the TEM. The stage is enclosed with shielding which enables a clean working environment within the instrument needed to support long automation runs. The system is cooled via an Autofill system enabling long run times and unattended operation over days.

#### Electron Column

The electron optical column is a NiCol column (ThermoFisher). The SEM comprises of a field emission gun assembly with Schottky-emitter, with a dual objective with both field-free magnetic and electrostatic lenses. The electron column provides detection in-lens using back scatter and secondary electrons.

#### Plasma ion column

The PFIB column can switch between three ion species (xenon, argon, and oxygen) switchable plasma ion source. These species have both chemical and physical differences which allow for a range of milling effects.

### Sputter Target

An ion sputtering device comprising a platinum mass is used to create thin layers of conductive platinum. Here, the primary ion source is used at high current to remove atoms of metal from a target allowing a dense vapor to flow over the sample. This renders samples conductive for imaging via SEM.

### Gas injection system

Organoplatinum is an organometallic compound which is used as a protective cap to protect the leading edge of the sample during milling. It enables lamella of sensitive samples to be protected by masking stray ions from affecting the delicate samples. This is typically condensed on the surface of the sample around a few micrometres thick as part of the preparation flow. The compound is evolved from a crucible contained within a delivery mechanism which generates vapor and delivers it via a needle to the chamber in small aliquots. The low temperature of the sample traps the vapour on contact and this aggregates over time to form the layer.

### Sputter rate measurements

The sputter rate was determined for each aperture commonly used during lamella preparation and on each beam by milling cross-sections with a controlled dose. Rates were determined at 30 kV.

The sample was plunge frozen yeast on a grid. To give sufficient sample thickness to mill ramps, cross-sections were generally made on the thickest regions of yeast clumps.

A schematic of the method for sputter rate determination is shown in Supplementary Fig. 3a. It is well established that FIB milling is sensitive to surface topography^19^. To reduce these effects the sample was first coated in GIS and platinum, as is the case for lamella preparation. A ramp was then milled into this to produce a smoother sample surface. The stage was then rotated and tilted so that now smooth sample surface of the ramp is presented perpendicular to the FIB column for cross-sectional milling. Each condition was repeated three times and an average taken.

For the angular dependence study, plunge frozen HeLa cells were used. In this case the stage was tilted after milling the ramps to introduce the required milling angle.

The milled depth was measured using the SEM. For perpendicular milling, the SEM imaging occurs at an angle relative to the trench (52°). The perpendicular depth was therefore calculated using the angle of imaging (i.e., tilt correction). The milled cuboid volume was calculated using this perpendicular depth and the known milling area, as shown in Equation 1.

For milling at non-normal angles, the milled volume was assumed to be a parallelepiped (side profile of this is shown in Supplementary Fig. 3e) with two rectangular and two square faces; the volume in this case is given in Equation 2. The volume was calculated by first calculating the area of the parallelogram (shown in yellow in Supplementary Fig. 3e) milled into the sample. The equations governing the relationship between the distances measured using the SEM and the milled volume dimensions are shown in Supplementary Fig. 3e.

Theta (*θ*) is the angle between the SEM and FIB column, in our case 52°, and alpha (*α*) is the milling angle relative to the milled ramp surface. Distances *bm* and *zm* were measured with the SEM without stage movement and trigonometry applied to work out the distances *b* and *z*. From *z* the distance *h* can be calculated and therefore area of the parallelogram subsequently. The volume was obtained by multiplying this calculated area by the width, *w*, of the initial milling square pattern.

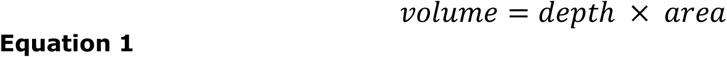

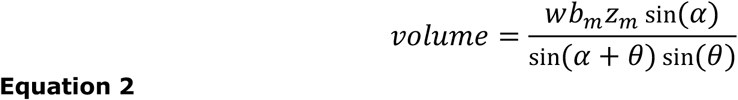

The sputter rate in μm^3^/s was calculated for each mill by dividing the calculated volume by the milling time. Sputter rates in μm^3^/nC were calculated by also dividing by the beam current that was measured for the chosen aperture.

Each mill was a standard square pattern with the Thermo Fisher Scientific default Silicon milling settings, which by default will set the pitch size and the dwell time. The pitch size was set as suggested as it is sample independent but milling time was changed.

Error bars shown on the plots in Supplementary Fig. 3 were calculated by taking the standard deviation of the 18 repeats (three repeats for six different times) at each beam current. Errors shown in Table 2 are the standard error derived using the least squares method from a linear line of best fit on the sputter rate (μm^3^/s) against beam current (nA) plots. The linear line of best fit was forced to pass through the origin. Error bars shown in Table 3 are the same as those on plots shown in Supplementary Fig. 3g and h and are the standard deviation.

### Cell culture

UltrAuFoil on gold 200 mesh R2/2 grids (Quantifoil) were subjected to micropatterning using the Primo module from Alveole mounted on a Leica DMi8 microscope following the manufacturer procedure. Briefly, the grids were coated with polylysine (100 μg/ml, 30 min) followed by mPEG-SVA (100 mg/ml, 1 h) and PLPP gel (1 h) prior to exposure to UV (50 mJ/mm2) to create circular patterns of 40 μm diameter. Then the grids were profusely rinsed with PBS before incubation with fibrinogen couple to Alexa 633 (Thermo Fisher). Micropatterned grids where then stored in Hank’s Balanced Salt Solution (HBSS) (Gibco).

HeLa cells were grown in Dulbecco’s Minimal Essential Media (DMEM) (Gibco) with high glucose and non-essential amino acids complemented with 10% FBS, 1% glutamine and gentamycin (25 μm/ml). Cells were seeded on micropattern grids for 2 h before washing. Infection with *C*.*trachomatis* LGV02 were performed as previously described^39^. 24 h post infection cells were PF using the Vitrobot (Thermo Fisher) offsetting the blotting pad to favour back blotting. Just prior the PF, the cells media has been replaced with the complete media with 10% glycerol (v/v). Vitrified grids were then clipped (Thermo Fisher) into autogrids (Thermo Fisher) and subsequently stored under liquid nitrogen.

Plunge frozen *S*.*cerevisiae* (Yeast) samples were prepared as previously described^38^.

### Automated lamella fabrication

Autogrid clipped TEM grids were loaded into a cassette and loaded into the PFIB’s multispecimen cassette which was then subsequently loaded onto the stage using the robotic sample delivery device (termed Autoloader) (Thermo Fisher Scientific). The SEM was used to screen the grids to ensure suitability for lamella preparation (Fig. 1a). Once selected for milling, grids were coated with trimethyl(methylcyclopentadienyl)platinum(IV) using the gas injection system (GIS) and then sputter coated with platinum metal using an inbuilt platinum mass which was targeted with the FIB beam at 16 kV and 1.4 µA to liberate clusters for adjacent surface metal coating.

Lamella sites were then identified and loaded into the AutoTEM cryo Software (Thermo Fisher Scientific) with the stage at eucentric height. The milling was then carried out using 30 kV argon automatically by the software. The milling procedure was as follows: eucentric height was refined at each site before milling stress relief cuts 5 µm each side of the intended lamella using a 2.0 nA ion beam. Three rough milling steps were then used to remove material above and below the intended lamella position: i.) at 2.0 nA, 0.74 nA and 0.2 nA. This left a lamella ~700 nm thick. This thicker lamella was then “polished” using 60 pA and 20 pA down to a nominal software target thickness of 70 to 90 nm. As the software is intended originally to operate with gallium, a considerable offset exists between the target thickness and the final lamella thickness for plasma ion sources. Rough milling was carried out first on all lamella sites before proceeding onto the final polishing step. Drift corrected milling was used to ensure accurate milling. Each pattern was the default Thermo Fisher Scientific ‘Rectangle’ with Silicon milling settings. All milling was done at 30 kV.

### Cryo-electron tomography

Multi-specimen cassettes with grids were directly transferred from the FIB/SEM to the transmission electron microscope via an Autoloader (Thermo Fisher Scientific). Tomography data were acquired with a Titan Krios (Thermo Fisher Scientific) transmission electron equipped with a Falcon 4 camera and a Selectris energy filter. Dose-symmetric tilt-series were collected using Tomo 5 (Thermo Fisher scientific) software at a nominal magnification of 64000 x magnification (corresponding to a calculated magnification of 81081 x) at a 1.85 Å pixel size in electron counting mode from 51° to −51°, corrected for the pre-tilt of the lamellae, with 3° increments, with a total dose of 175 e^−^/Å^2^.

### Sub volume averaging

Warp version 1.0.9^40^ was used for gain- and motion correction using 10 frame groups per tilt, contrast transfer function (CTF) correction and creating tilt-series stacks. Tilt-series were aligned and reconstructed at a binning factor 8 using AreTomo 1.1.1^41^ and tomograms were flipped using IMOD 4.11^42^, followed by bandpass filtering using EMAN 2.91^43^. Ribosomes were automatically picked using crYOLO 1.8.3^44^ by creating a training dataset from 5 representative tomograms, followed by prediction on 180 tomograms, identifying 70436 putative ribosomes. Particles were extracted at a binning factor 4 using Warp with a box size of 64×64×64 pixels and imported into RELION 3.1^45^. An initial reference was generated by performing 3D refinement on a random subset of 1000 particles, followed by 3D refinement with all the particles. 3D classification was performed with a circular mask, identifying 18119 particles as ribosomes. Another round of 3D classification with a tighter ribosome-shaped mask was performed, yielding 8 classes with 80S ribosomes totalling 16204 particles, one class of the 60S ribosome with 1896 particles and one junk-class with 19 particles. The 16204 particles from 80S ribosome classes were extracted with warp at binning 2 with a box size of 128×128×128 pixels. Particle poses were subjected to 3D refinement in RELION using a thresholded ribosome mask. Particles poses for 15628 particles (six tomograms could not be refined post-RELION in M due to program errors) were imported into M version 1.0.9^33^ for 5 rounds of refinement where sequentially the following additional parameters were solved for: 1) 3 × 3 image warp grid and particle poses 2) 4 × 4 image warp grid 3) 3 × 3 × 2 × 10 volume warp grid 4) same settings as round 3 5) stage angles and per-particle defocus estimation. Structures determined were fitted using a human ribosome model (PDB: 4UG0^46^), and the pixel size adjusted empirically to fit the PDB model. The determined pixel size for the map was 1.9 Å. This value was used for all FSC calculations (0.143) and B-factor calculations.

To produce structures of ribosomes at different distances from the lamella edge, a custom Python script was written that was able to calculate the distance of particles from XYZ coordinates within the output from M/RELION. The software is freely available on GitHub: https://github.com/rosalindfranklininstitute/RiboDist. Briefly, boundary models were created manually for each tomogram every ~100 YZ slices using IMOD^42^ and converted into a text files for use in the script. The script interpolates the manually created boundary models, determines the nearest distance for each ribosome coordinate to the edge model, determines the distance of the top and bottom edge model in the centre of the tomogram and generates STAR format files with additional columns for the distance for every particle and the thickness of the lamella for each tilt-series. The column of values can then be used as threshold for generating STAR files in RELION3.1^45^. All particles with distances less than 30 nm were then parsed into a new STAR file and the number of particles used to generate a second randomised STAR file from the remaining particle list. The two STAR files were then used for new rounds of sub-volume averaging in RELION3.1. The local resolution was then determined after import into M, without any further refinement.

B-factors were calculated using RELION based on Rosenthal and Henderson^34^. Briefly, reconstructions were determined for subset of the particles from within the 30 nm layer and beyond this distance. The determined resolutions were then plotted as a function of the particle number and the 2 over slope of the linear fit calculated (Fig. 4).

### Data analysis

To determine the distance where tilt-series were recorded to the platinum layer in the front of the lamellae, search images saved by Tomo 5 were stitched in FIJI^47^ and the distance from each position to the front of the platinum layer along the direction of the FIB beam was measured with the line measurement tool.

## Acknowledgements

The authors would like to thank Lu Gan, Alex de Marco, and Sebastian Tacke for robust discussions around sputtering rates. This work was supported by the Wellcome Trust through the Electrifying Life Science project (220526/Z/20/Z to J.H.N). The Rosalind Franklin Institute is funded by UK Research and Innovation through the Engineering and Physical Sciences Research Council (EPSRC).

## Author Contributions

M.D. prepared samples for cryo-FIB. C.B., M.D., and T.G. performed cryo-FIB, and collected cryoET data. C.B. and M.G. performed sub-volume averaging. N.B.y.Y. implemented new computation tools to help with data analysis. J.M.M., and Z.P., managed the development of the prototype cryo-FIB/SEM instrument, with input from M.D., J.H.N. and M.G. C.B., T.G., and M.G. prepared figures. J.H.N. and M.G. supervised the project. C.B. and M.G. wrote the manuscript. All authors reviewed the data and commented on the manuscript.

## Supplementary Information

The online version contains supplementary material available at XX

## Data Availability

Sub volume averages are deposited in the Electron Microscopy Data Bank (EMDB) under the following codes: EMD-XXXX (full reconstruction), EMD-XXXX (lamella edge proximal) and EMD-XXXX (away from lamella edge). The frames and associated metadata needed to reconstruct the data (180 tomograms) have been deposited on the EMPIAR data server (EMPIAR-XXXXX).

## Supplementary Figures

**Supplementary Figure 1.**
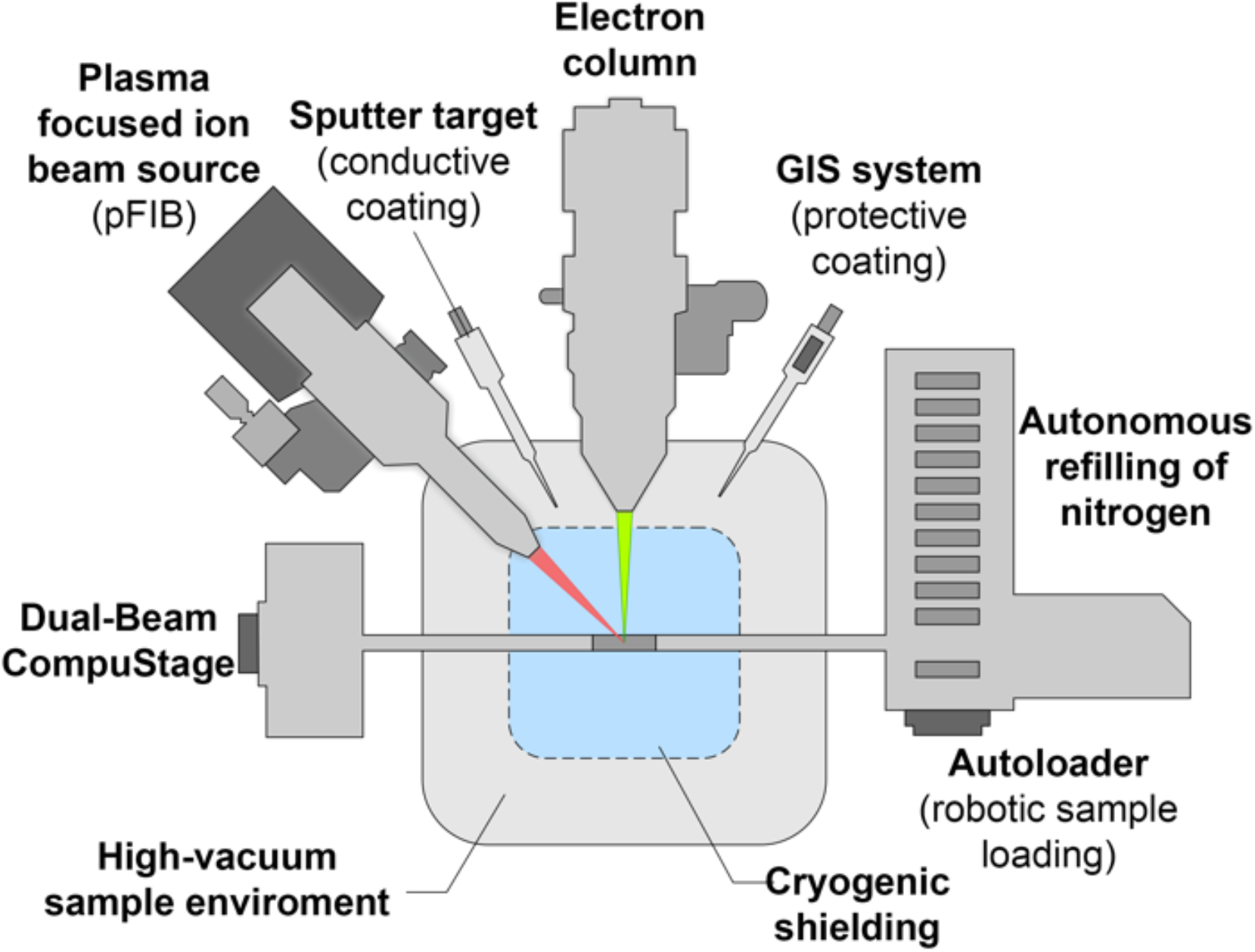
Schematic overview of the FIB/SEM microscope used in this study. The microscope has several elements that make it amenable to lamella production on long time scales in an automated fashion. This includes the high-vacuum chamber (10^−7^ mbar), multi-specimen loading and automatic refilling with liquid nitrogen. The instrument is equipped with a Hydra-type focused ion beam column, which utilises xenon, argon, or oxygen gas sources for plasma generation. The CompuStage enables robust position of the sample within the co-incidence point with ~500 nm accuracy. Gas injection system (GIS) and sputter target enable protective layer and charge mitigation functionality to enable scanning electron imaging using the NiCol SEM column.

**Supplementary Figure 2.**
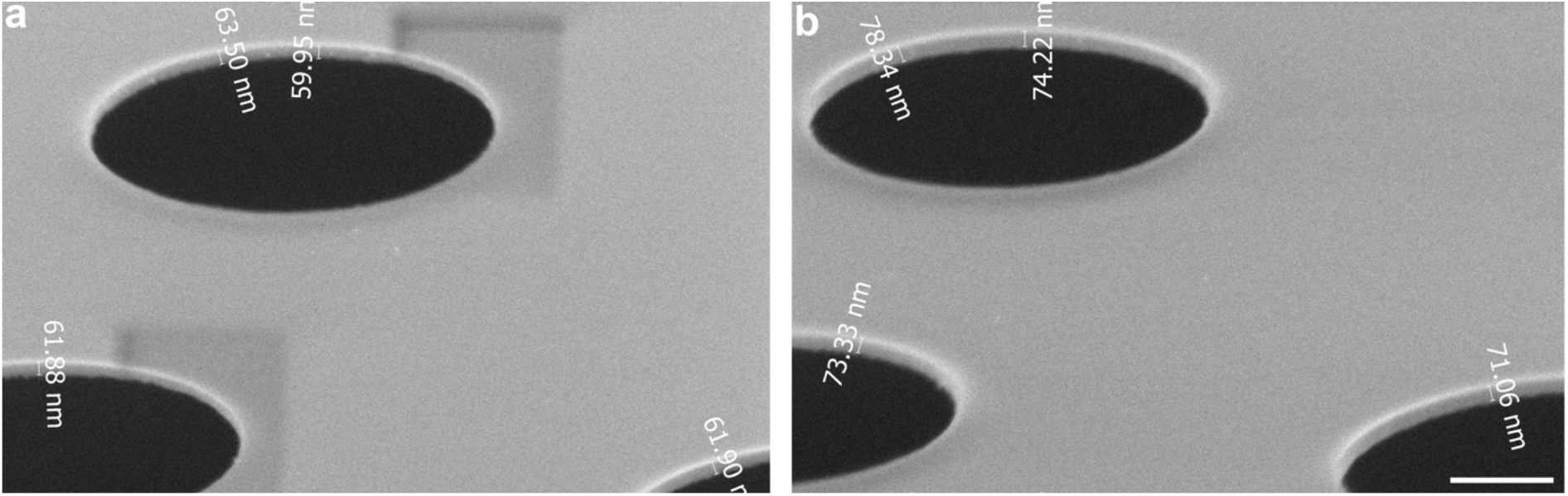
Ice growth over time in in the column of the FIB/SEM microscope. SEM images of the same location of a TEM grid were acquired at 0 h (a) and 8 h (b) and the EM grid foil thickness measured before and after, corrected for the stage tilt. This gave an average ice growth rate in the column of 1.6 nm/h. Scale bar: 500 nm. The pixel size for these images was 1.3 nm.

**Supplementary Figure 3.**
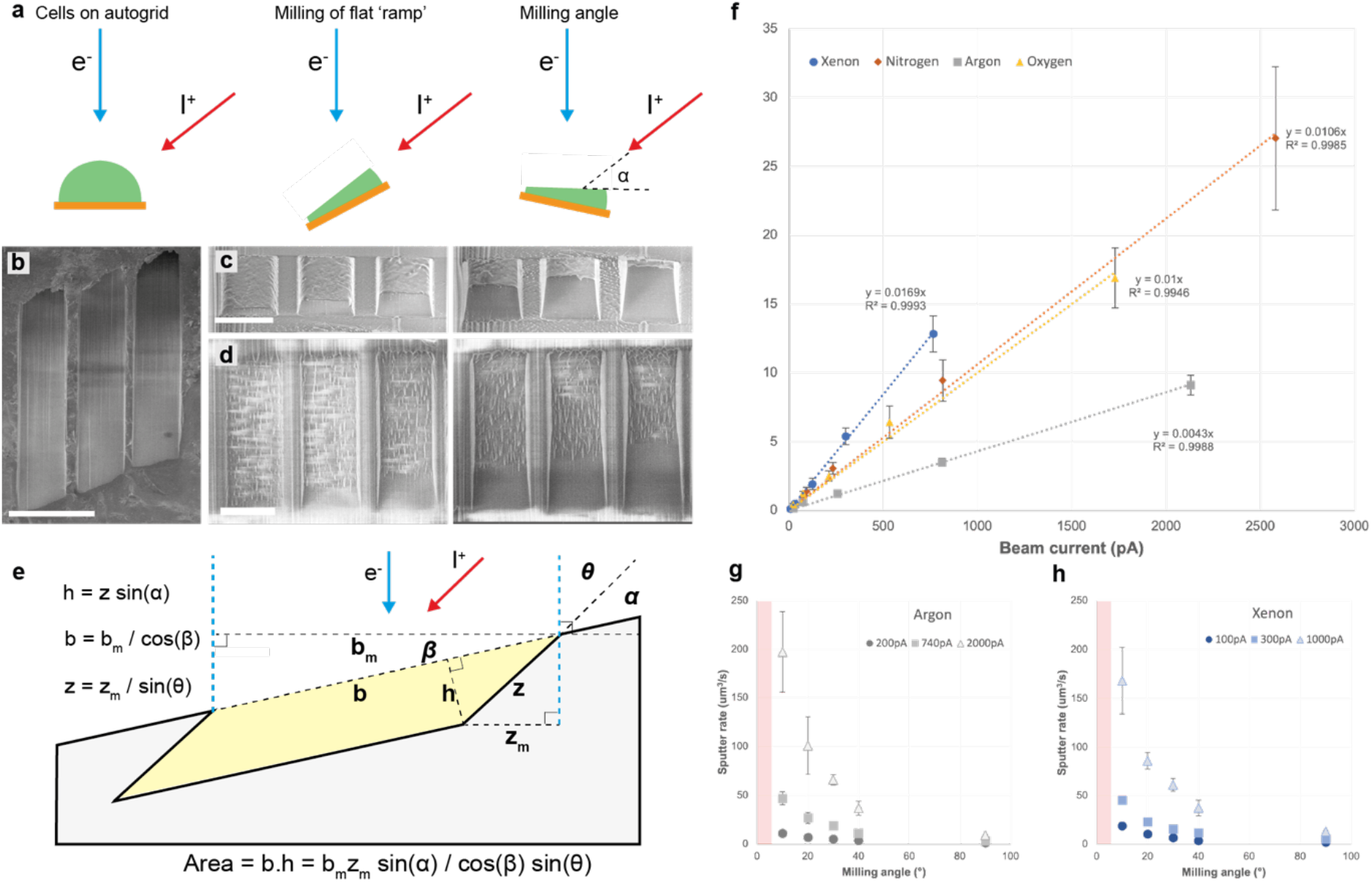
Determination of PFIB sputter rates at 30 kV on vitrified yeast samples. **(a)** Cartoon showing sputter rate measurement process, which involves milling a flat ramp **(b)** before tilting to present this fresh surface to the FIB at a milling angle of alpha. Six rectangles of varying exposure time were then milled and repeated three times, giving 18 measurements for each condition. Representative images of milling angles of 90° and 20° are shown in **(c)** and **(d)** respectively. The depth of the milled trench increases in **(c)** and **(d)** from left to right, as a function of milling time. The volume was then calculated from measurement of the milled depth using the SEM, as shown in **(e)** from which the sputter rate could be calculated (see materials and methods). **(f)** Sputter rate vs current for xenon, nitrogen, argon, and oxygen at a range of currents with a milling angle of 90°. Error bars are the standard deviation. **(g)** and **(h)** show the sputter rate as a function of milling angle for three different beam currents of argon and xenon respectively. The red area shows where the trend is expected to reduce back to zero but was not included as accurate measurement of such low angle mills was experimentally challenging. Error bars are the standard deviation.

**Supplementary Figure 4.**
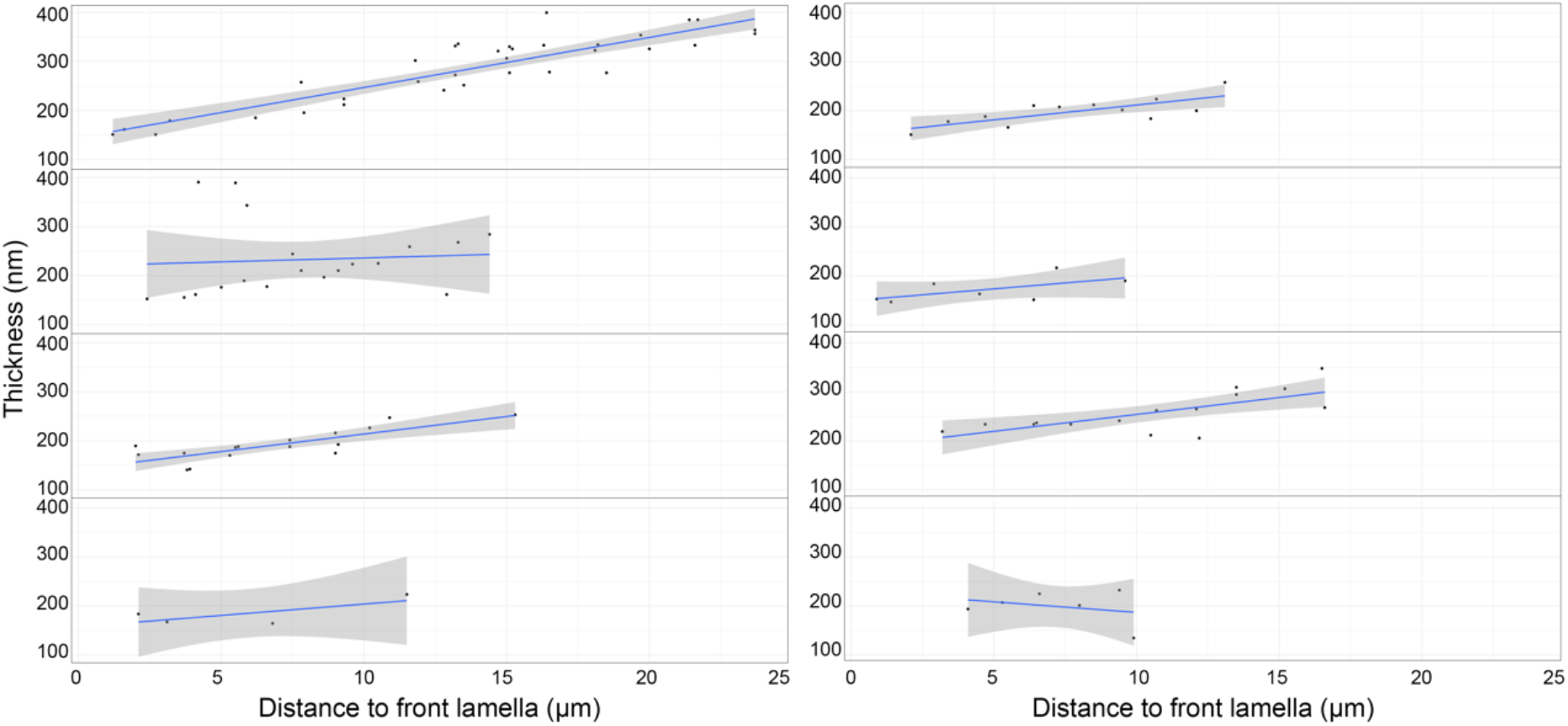
Distances from the edge of the front of the lamella for tomograms acquired for 8 lamellae vs the measured thickness for a given tomogram. Plots are shown for 8 lamellae of the distribution of thickness against the distance form the leading edge of the lamella. The plots show the approximate thickness profile of the lamella and indicate their flatness.

**Supplementary Figure 5.**
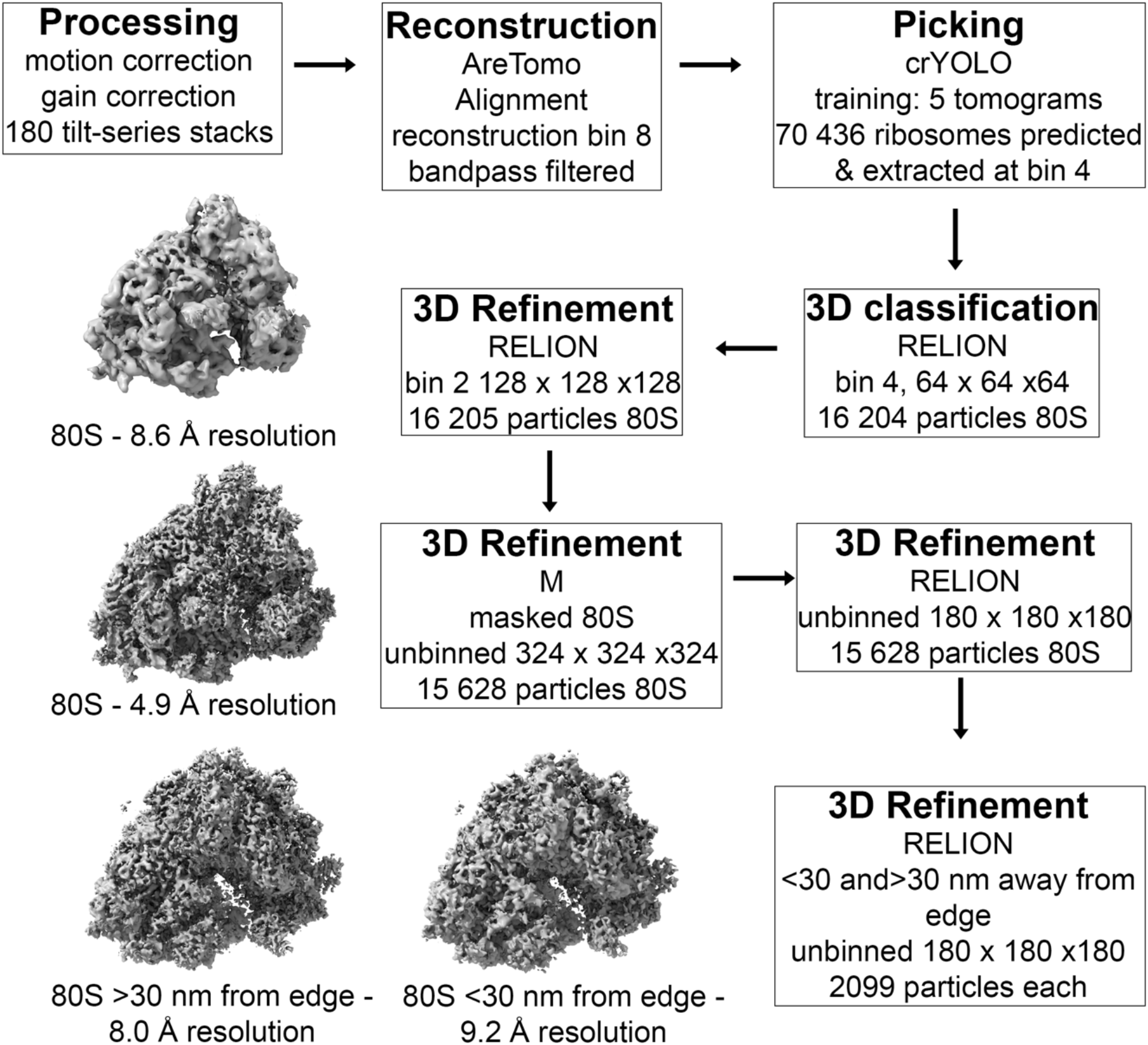
Schematic overview of sub volume averaging workflow. All 180 tilt-series were processed, reconstructed and putative ribosomes were identified with crYOLO. Further 3D classification in RELION and subsequent 3D refinement in RELION and M yielded a 4.9 Å reconstruction of the 80S ribosome, and two separate reconstructions with a subset of the particles based on their distance from the lamella milling edge.

**Supplementary Figure 6.**
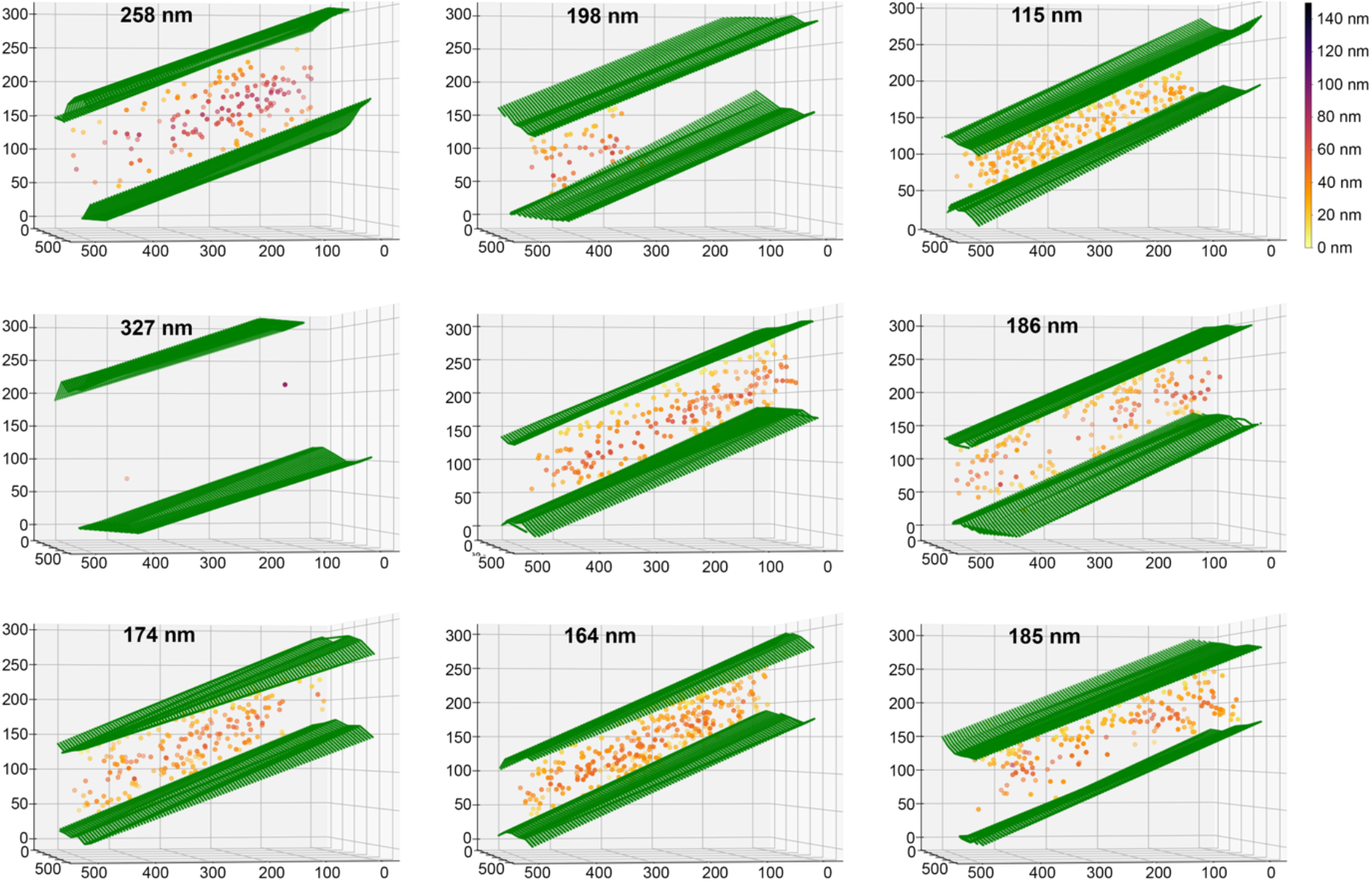
Distribution of ribosome location within 9 representative tomograms. Graphical representation of the annotated lamella edges (green) of 9 different tomograms. Ribosomes used for sub-volume averaging (see Fig. 3) are shown, where the colour of the ribosome points are indicative of their distance from the edge (in nanometres). XYZ coordinates are indicated for 8 x down sampled tomograms (pixel size: 1.48 nm).

**Supplementary Figure 7.**
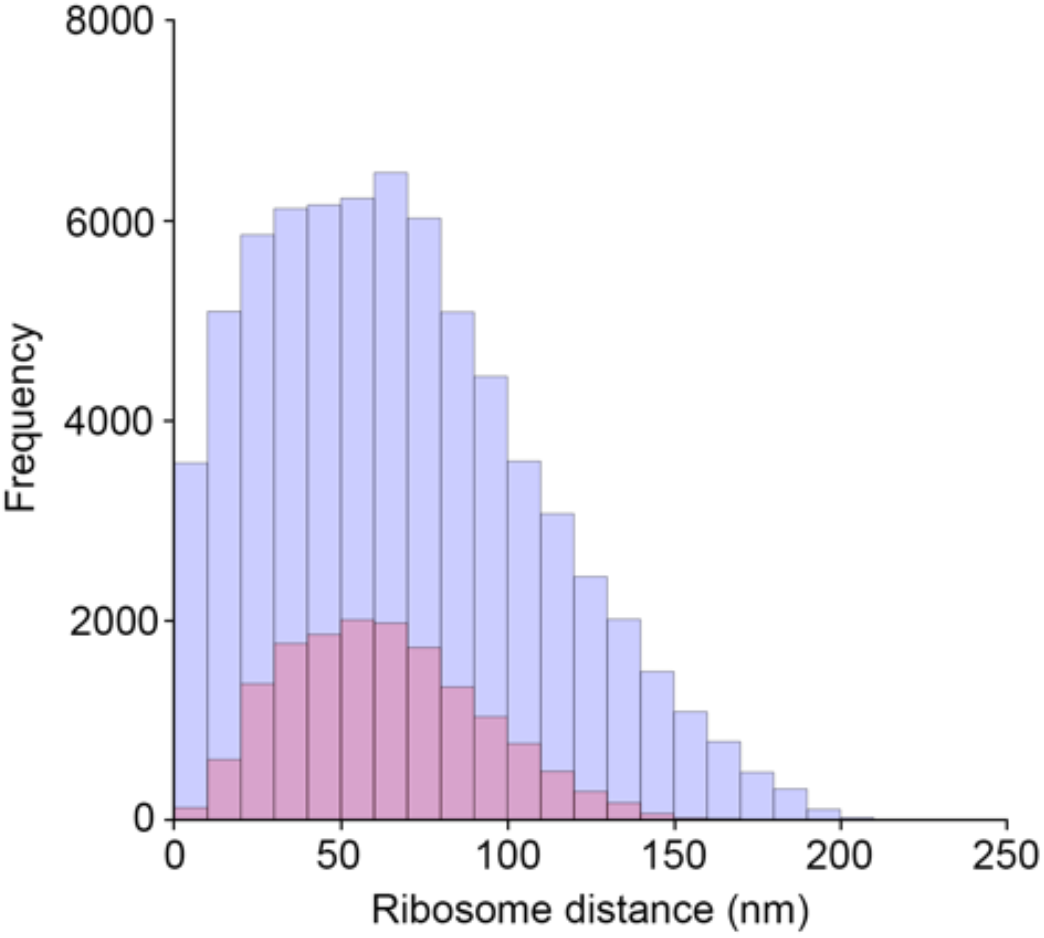
Distribution of ribosome distances to the milling edge before and after 3D classification. The frequency putative-ribosomes identified with crYOLO (blue) and 80S ribosomes after 3D classification (red) based on the ribosome distance in nanometers (bin size: 10 nm).

**Supplementary Figure 8.**
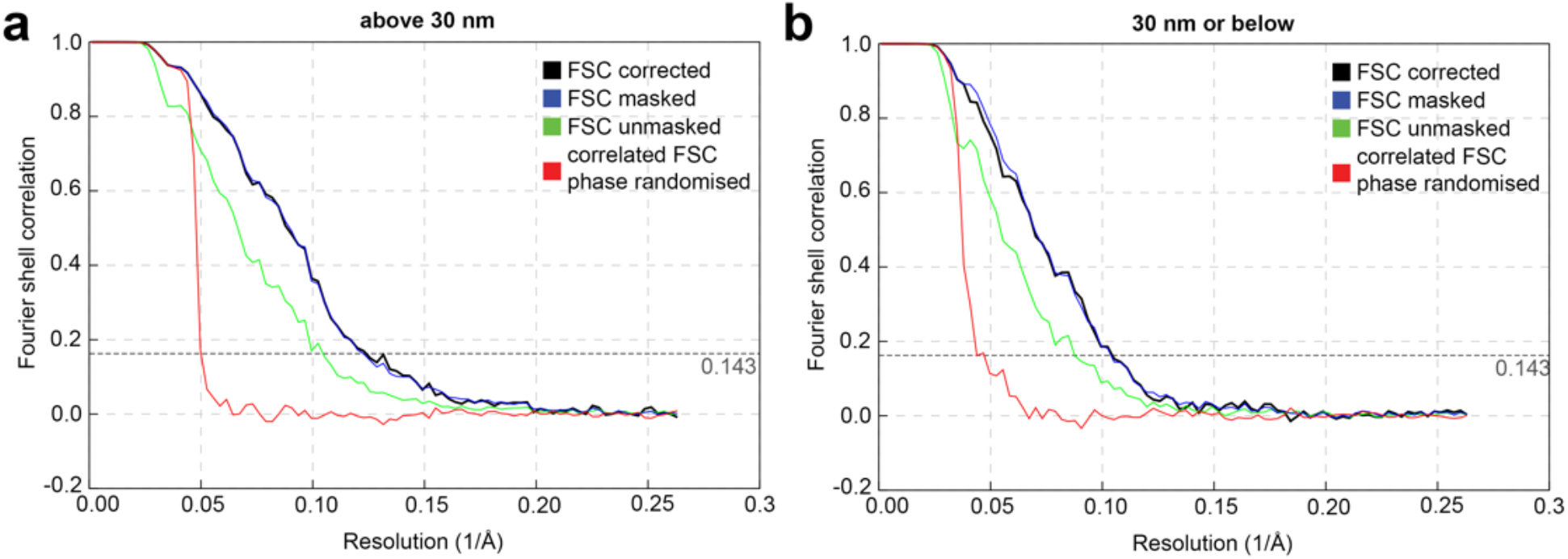
FSC curves for ribosome structures from within 30 nm distance from lamella edge and in the centre. FSC curves are shown for ribosomes structures determined from: (a) 2099 ribosomes greater than 30 nm from the lamella edge and (b) 2099 ribosomes within 30 nm of the lamella edge. Resolutions are 8.0 Å and 9.2 Å, respectively.

## Supplementary Videos

**Supplementary Video 1**. Tomographic volume of the slice shown in Fig. 1h. Scalebar: 100 nm.

**Supplementary Video 2**. Video of the 3-dimensional volume local resolution map shown in Fig. 3b. Scale bar is 10 nm

**Supplementary Video 3**. Sub volume average density map of the ribosomal L4 subunit obtained in this study overlaid with the previously obtained structure of isolated human ribosomes (PDB: 4UG0^46^). Scale bar is 1 nm.

